# Muscle Energy Crisis is the Initial Driver of Locomotor Dysfunction in β-glucuronidase-Deficient *Drosophila*

**DOI:** 10.1101/2025.10.25.684497

**Authors:** Nishan Mandal, Soudipa Bhattacharjee, Rupak Datta

**Author notes:** Author for correspondence: Rupak Datta; Extn: 1214.

## Abstract

Mucopolysaccharidosis type VII (MPS VII) is characterised by progressive locomotor decline, attributed to musculoskeletal and neurological defects. However, the severity varies according to the extent of β-glucuronidase (β-GUS) deficiency, with musculoskeletal deformities typically preceding neurological manifestations. To investigate the underlying cause of neuromuscular pathology, we employed β-GUS-deficient *Drosophila* models. In *Drosophila*, β-GUS is encoded by two genes, CG2135 and CG15117. Previously generated CG2135^-/-^ flies recapitulated several hallmark features of MPS VII, despite retaining approximately 30% residual β-GUS activity contributed by CG15117. To assess the consequences of complete loss of β-GUS, we now generated CG15117^-/-^ flies using CRISPR/Cas9 and subsequently established CG15117^-/-^;CG2135^-/-^ double knockout (DKO) flies. Phenotypic assessment revealed differences in susceptibility to starvation, lifespan, and locomotor function, with DKO flies exhibiting more severe impairments than either single knockout flies. Notably, CG15117^-/-^ did not show any significant defect in lifespan or locomotion, except under starvation conditions. Consistent with our earlier finding in CG2135^-/-^ fly brains, ATP depletion in DKO fly brains became evident only after 45 days of age, failing to explain the locomotory defects observed in earlier ages. Interestingly, analysis of muscles of 30-day-old CG2135^-/-^ and DKO flies revealed abnormal mitochondrial accumulation with autophagy defect and severe ATP depletion. Consequently, these defects led to locomotor impairment driven by apoptotic muscle degeneration. Collectively, this work provides the first evidence of tissue-specific vulnerability in MPS VII models, identifying muscle as an early pathological target and offering new insights into disease progression.

## Introduction

Mucopolysaccharidosis is a progressive, multisystem disorder caused by the lack of β-glucuronidase, a lysosomal hydrolase involved in the degradation of glycosaminoglycans (GAGs) (Shipley et al., 1993; Sly et al., 1973). Progressive skeletal deformities, hydrops fetalis, growth impairment, organomegaly, visual impairment, recurrent infections, and a range of neurological anomalies are common clinical characteristics associated with MPS VII. Lysosomal GAG storage in different cell types causes multisystem failure, ultimately leading to premature death. Musculoskeletal and neurological abnormalities associated with MPS VII patients are commonly attributed to impaired mobility (Montaño, Lock-Hock, et al., 2016; Shipley et al., 1993). Progressive loss of muscle strength, increased musculoskeletal pain, and stiffness, along with cardiac myopathy in MPS VII patients, indicate muscular atrophy (Jones et al., 2021; Montaño, Lock-hock, et al., 2016; Oldham et al., 2022). Similarly, impaired cognition, speech delay, and hearing loss are cardinal features of neurological defects observed in MPS VII patients (Montaño, Lock-hock, et al., 2016). However, the severity of these clinical manifestations could vary depending on the mutation acquired in the β-GUS gene. It is noteworthy that in milder disease forms, neuropathological defects were either completely absent or appeared only at advanced stages, unlike musculoskeletal abnormalities, suggesting differential tissue vulnerability (De Kremer et al., 1992; Montaño, Lock-hock, et al., 2016; Storch et al., 2003). While enzyme replacement therapy (ERT) offers partial benefit, its restricted delivery to tissues like the brain and muscle highlights the necessity for improved understanding to design alternative interventions (Giugliani et al., 2024; McCafferty & Scott, 2019; Sahin et al., 2022). However, the absence of detailed mechanistic understanding constrains the development of rational, targeted therapeutic approaches.

To elucidate the molecular mechanism underlying the progression of MPS VII, we employed β-GUS- deficient *Drosophila melanogaster* as a model system. In *Drosophila*, two genes annotated as *CG2135* and *CG1117* encode functional β-GUS enzymes. In vitro enzymatic assays demonstrated that CG2135 possesses approximately six-fold higher activity than CG15117. Based on this observation, the CG2135 gene was knocked out to generate the CG2135^-/-^ fly strain. Although these flies retained approximately 30% residual β-GUS activity due to the presence of CG15117, they nonetheless recapitulated several cardinal features of MPS VII patients. Among the disease-related phenotypes, a progressive reduction in climbing ability was particularly evident in these flies after 21 days of age, which further worsened with age. This progressive decline in locomotor performance observed in CG2135^-/-^ flies is indicative of neuromuscular degeneration (Bar et al., 2018). We have recently reported that impaired mitophagy in the brains of these flies leads to the accumulation of dysfunctional mitochondria, culminating in apoptotic neurodegeneration (Mandal et al., 2025). However, such mitochondrial defects in the brain became apparent only at later stages (at 45 days of age) of disease progression and therefore cannot account for the locomotor decline observed at earlier stages. Furthermore, it remains unclear whether loss of CG15117 alone, or simultaneous disruption of both β-GUS-encoding genes, would exacerbate disease-associated phenotypes.

In addition to our findings in the CG2135^-/-^ fly brain, mitochondrial dysfunction and altered energy metabolism have also been reported in the brains of MPS IIIC mouse models (Martins et al., 2015; Settembre et al., 2008). Although limited, these studies collectively support the involvement of mitochondrial dysfunction in the neuropathology of mucopolysaccharidoses. However, comparable investigations in muscle tissue are notably lacking. Given that high-energy-demanding tissues such as the brain and muscle rely critically on efficient mitochondrial quality control that involves the autophagy-lysosome axis, defects in these processes are likely to be particularly detrimental (Lei et al., 2024; Shen et al., 2022). It is therefore equally imperative to examine mitochondrial status within muscle tissue as in the brain to better understand its contribution to disease progression.

In this study, we investigated the impact of partial versus complete loss of β-GUS activity on disease progression, with a particular focus on muscle degeneration. Using CRISPR/Cas9, we generated a CG15117 knockout strain, which was subsequently brought into the background of CG2135^-/-^ flies to obtain a double knockout line. Complete loss of β-GUS activity markedly worsened the disease-related phenotypes in comparison to CG2135 knockout flies, as reflected by a shorter lifespan and more severe locomotor decline. By contrast, CG15117^-/-^ flies displayed no obvious phenotypes other than increased sensitivity to total starvation. Our previous work established that locomotor decline in CG2135^-/-^ flies was driven by extensive clearance defects, mitochondrial dysfunction, and consequent ATP depletion in the brain (Mandal et al., 2025). Although consistent with our previous observation in CG2135^-/-^ fly brain, depleted brain ATP level in DKO flies was found to be exclusively restricted to the late stage of the disease (45 days of age). This temporal discrepancy failed to account for the locomotor impairment observed in these flies at earlier stages. However, as locomotion relies not only on the brain but also on muscle function, we extended our analysis to assess mitochondrial status in muscle tissue. Our study revealed a significant ATP depletion in the muscles of 30-day-old CG2135^-/-^ and DKO flies. ATP loss was previously recorded in 45-day-old CG2135-/- fly brain as a result of defective mitophagy (Mandal et al., 2025). A similar abnormal accumulation of mitochondria along with defective autophagy-lysosome clearance pathways in 30-day-old CG2135-/- and DKO fly muscles indicates that defective mitochondrial clearance could be the plausible cause of ATP depletion. This energy crisis finally led to apoptotic muscle degeneration and locomotor defects. This study provides evidence that muscle is affected prior to neuronal tissues due to loss of β-GUS activity. Our results thus report tissue-specific temporal variation in pathological outcome due to β-GUS-deficiency and muscle degeneration as the initial driver of locomotor defect.

## Materials and Methods

All reagents were purchased from Sigma-Aldrich unless specified otherwise. Information on oligonucleotides and antibodies used in this study is provided in Supplementary Tables 1 and 2, respectively.

### Fly Culture

All fly stocks are raised and maintained in standard cornmeal food. To ensure the reproducibility of the experimental result, all the flies were grown at a controlled temperature of 25°C and a 12-hour light-dark cycle to simulate day and night conditions. For the aging experiments, newly eclosed flies were collected and maintained till they attained experimental age. These flies were transferred to fresh food vials at an interval of two days. All fly stocks used in this study were obtained from Bloomington Drosophila Stock Centre at Indiana University, USA W^1118^, TM3Ubx/TM6Be, Tft/CyO, Nos-Cas9 (54591). CG2135^-/-^ flies were previously generated by our lab (Bar et al., 2018). pCFD4-CG15117-sgRNA, CG15117^-/-^ and CG15117^-/-^;CG2135^-/-^ fly strains were generated in this study.

### Generation of CG15117^-/-^ and CG15117^-/-^;CG2135^-/-^ double knockout (DKO) fly strains

To generate a CG15117 knockout Drosophila strain, the CRISPR/Cas9 system was employed using two sgRNAs targeting the 5′ and 3′ ends of the CG15117 gene to induce a large genomic deletion (Port et al., 2014). The sgRNAs were designed using the CHOPCHOP server and validated for specificity using BLAST (Labun et al., 2019). These were cloned into the pCFD4-U6:1_U6:3 tandem sgRNA expression vector via Gibson assembly. The sgRNA core of the pCFD4 plasmid was PCR-amplified using forward and reverse primers containing the gRNA1 sequence and the reverse-complemented gRNA2 sequence, flanked by promoter and gRNA core sequences, respectively. The amplicon and BbsI-digested pCFD4 vector backbone were extracted following agarose gel electrophoresis and subsequently ligated using the Gibson assembly kit following the manufacturer’s protocol. The recombinant plasmid was transformed into E. coli (DH5α), screened via colony PCR using gRNA1 as the forward primer and the reverse-complemented gRNA2 as the reverse primer, and verified by sequencing. The verified plasmid was injected into Drosophila embryos to generate sgRNA-expressing flies, which were then crossed with Nos-Cas9 flies for germline editing. A detailed crossing scheme is provided in the Supplementary Fig. 1. CG15117 knockout progeny were screened based on genetic markers and confirmed via PCR. To generate a double knockout strain (CG15117^-/-^;CG2135^-/-^), multiple genetic crosses were performed using the single knockout lines (Fig. S2). The resulting DKO flies were validated by PCR using gene-specific primers.

### Lifespan Assay

To assess the longevity of β-GUS-deficient and wild-type flies, a lifespan assay was performed. Synchronized 1-day-old male and female flies were collected and maintained together for 4 days to allow sexual maturation. On day 4, flies were segregated by sex into cohorts of 20 individuals per genotype. Flies were maintained under standard conditions and transferred to fresh food vials every other day. The number of dead flies was recorded daily until all individuals had perished. Survival data were used to generate Kaplan-Meier survival curves.

The log-rank test was applied to compare survival distributions between genotypes. Mean lifespan was calculated following Lushchak et al.,2012 (Lushchak et al., 2012).

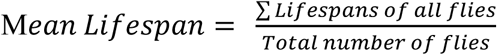

### Starvation Assay

To evaluate the sensitivity of flies to nutrient deprivation, a starvation assay was conducted on 30-day-old male flies. Flies were initially collected at day 1 post-eclosion and maintained under standard conditions until 30 days of age, with regular transfer to fresh food vials every other day. For the assay, 30-day-old males were randomly grouped into cohorts of 10 flies and transferred into empty vials containing a piece of water-soaked tissue to provide moisture without nutrition. The number of deceased flies in each vial was recorded at 12-hour intervals until all flies had perished. Mean survivability time was calculated using the same formula employed in the lifespan assay (Lushchak et al., 2012). This assay was used to assess starvation sensitivity in β-GUS-deficient strains relative to wild-type controls.

### Climbing Assay

To assess locomotor activity, a climbing assay was performed on β-GUS-deficient fly strains and wild-type controls (Bar et al., 2018). Age-matched flies were collected on day 1 post-eclosion and maintained under standard conditions until the respective assay time points. For each genotype, cohorts of 20 male flies were prepared one day prior to the experiment. The climbing index was defined as the percentage of flies that successfully crossed the 25 mL mark within 15 seconds. Climbing assays were conducted at 7, 15, 21, 30, and 45 days of age. Age-matched wild-type (WT) flies served as wild-type controls. Statistical significance was assessed using multiple t-tests followed by the Holm-Sidak error correction method.

### β-GUS activity assay

β-Glucuronidase (β-GUS) activity was measured fluorometrically using 4-methylumbelliferyl β-D-glucuronide (4-MUG) as the substrate (Glaser & Sly, 1973). Flies were homogenized in 1X-PBS containing protease inhibitor cocktail (PIC) using plastic pestles, followed by sonication at 100% amplitude for three cycles (15-second pulse with 10-second rest) on ice. The lysate was centrifuged at 14,000g for 15 minutes, and the supernatant was collected for analysis. For the assay, 10 μL of supernatant was incubated with 100 μL of 10 mM 4-MUG substrate solution at 37 °C for 1 hour. The reaction was terminated by adding 1.9 mL of glacial carbonate stop buffer (pH 10.5), and fluorescence was measured using a fluorimeter with excitation/emission settings of 365/460 nm. A standard curve was generated using increasing concentrations of 4-methylumbelliferone (4-MU) in glacial acetic acid. The linear range of the standard curve was used to determine the amount of hydrolyzed substrate. Total protein concentration was quantified by the Lowry method using BSA as a standard. β-GUS activity was expressed as nanomoles of substrate hydrolyzed per hour per milligram of protein (units/mg).

### Genomic DNA isolation

Genomic DNA was isolated from 10-15 adult flies using the phenol-chloroform extraction method. Flies were lysed in 400 μL of lysis buffer (100 mM Tris-Cl pH 7.5, 100 mM EDTA, 100 mM NaCl, 0.5% SDS) and incubated at 65 °C for 30 minutes. Subsequently, 800 μL of Buffer B (prepared from 5 M potassium acetate and 6 M lithium chloride) was added and samples were incubated at room temperature for 1 hour. After centrifugation at 14,000g for 15 minutes, 1 mL of the supernatant was collected and mixed with an equal volume of phenol:chloroform (1:1). The mixture was centrifuged again at 14,000g for 15 minutes at 4 °C. The resulting aqueous phase was transferred to a new tube and mixed with 600 μL of isopropanol, followed by a 10-minute incubation at room temperature. DNA was pelleted by centrifugation (14,000g, 15 minutes, 4 °C), washed with 75% ethanol, air-dried, and resuspended in nuclease-free water. The integrity of the isolated genomic DNA was confirmed by electrophoresis on a 1% agarose gel.

### Genomic DNA Isolation, RNA Isolation, and cDNA Synthesis

Total RNA was isolated from 10-15 fly heads using TRIzol reagent, following the manufacturer’s instructions. RNA integrity was assessed by electrophoresis on a 1.8% agarose gel prepared in 1X-TAE buffer. To eliminate genomic DNA contamination, 1 μg of RNA was treated with DNase I as per the manufacturer’s protocol. cDNA was synthesized using a commercial cDNA synthesis kit and oligo(dT) primers. Successful reverse transcription was confirmed by PCR amplification of the housekeeping gene RP49 using gene-specific primers.

### Western blotting

Western blotting was performed to analyze protein levels in β-GUS-deficient fly tissues. For sample preparation, 5-10 fly heads or muscle tissues were homogenized in 1X-Laemmli buffer supplemented with protease inhibitor cocktail and heated at 100 °C for 5 minutes. Equal amounts of protein were loaded onto SDS-polyacrylamide gels and separated by electrophoresis. Gels were equilibrated in transfer buffer and proteins were transferred to PVDF membranes using a semi-dry transfer system. Membranes were blocked with 5% skimmed milk in 0.025% TBST for 2 hours, washed, and incubated with primary antibodies overnight at 4 °C. Following washes, membranes were incubated with HRP-conjugated secondary antibodies for 1.5 hours at room temperature. Signals were developed using the Invitrogen SuperSignal chemiluminescent substrate and imaged with a Bio-Rad ChemiDoc system. Band intensities were quantified using ImageJ software and normalized to tubulin.

### ATP Quantification

Total ATP levels were measured using the ATP Determination Kit (Thermo Fisher Scientific), following the manufacturer’s instructions. Briefly, ten dissected fly heads or muscle tissues were homogenized in cold lysis buffer consisting of 1X-PBS supplemented with 1X-protease inhibitor cocktail (PIC). The homogenates were centrifuged at 20,000g for 20 minutes at 4 °C, and the supernatants were collected for ATP measurement. For the assay, 2 μL of each sample was added to 98 μL of ATP reaction solution and incubated for 5 minutes at 28 °C. Luminescence was recorded using a microplate reader. A standard curve generated using known concentrations of ATP was used to calculate the total ATP content in each sample. ATP levels were normalized to total protein content, which was determined using the Lowry method.

### TUNEL staining

Apoptotic cell death in β-GUS-deficient fly muscles was assessed using the TUNEL assay as described by Denton et al. (2008) (Denton et al., 2008). Briefly, 30-day-old fly muscles were dissected in S2 media supplemented with 10% FBS and fixed in 4% paraformaldehyde (PFA) containing 0.1% Triton X-100 for 30 minutes. Fixed samples were washed twice each with 0.1% and 0.5% PBS-Triton (PT) for 3 minutes, followed by permeabilisation in sodium citrate at 65°C for 30 minutes. After three additional washes in 0.5% PT, samples were incubated with the TUNEL reaction mixture (Roche) at 37°C for 3 hours, according to the manufacturer’s instructions. Samples were subsequently washed three times in 0.1% PT, counterstained with DAPI, and mounted in Vectashield. Imaging was performed the following day using Leica SP8 confocal microscope. TUNEL-positive puncta were quantified and represented as the number of TUNEL-positive nuclei per unit area.

### MitoTracker Staining

MitoTracker staining was performed on age-matched fly brains as described previously. Brains were dissected in S2 medium and incubated with 500 nM MitoTracker Red (Invitrogen) and 2 µg/mL Hoechst for 1 hour in the dark. Samples were then washed twice with 1X PBS and imaged immediately after mounting with S2 media using Leica SP8 confocal microscope. MitoTracker fluorescence intensity per 1000 X 1000 pixel unit area was quantified using ImageJ software.

### Immunostaining and Imaging

For immunostaining, fly muscles were dissected in S2 medium supplemented with 10% FBS (Avantor), 100 U/mL penicillin (Gibco), and 100 µg/mL streptomycin (Gibco). Samples were fixed in 4% paraformaldehyde (PFA) for 30 minutes at room temperature, washed in 0.3% PT (1× PBS with 0.3% Triton X-100), and blocked in PBT × GS (0.3% PT with 0.5% BSA and 5% normal goat serum) for 2 hours. Samples were incubated with primary antibodies for 36 hours at 4°C, washed five times in 0.3% PT, and then incubated with secondary antibodies for 12 hours at 4°C. After five further washes, samples were counterstained with DAPI (1 µg/mL) and mounted in Vectashield (Vector Laboratories). Imaging was performed using a Leica SP8 confocal microscope. Image processing and quantification were carried out using LasX and ImageJ software. For mean intensity measurements, fluorescence intensity was calculated from multiple ROIs per field using ImageJ.

### Statistical analysis

All the experiments were done at least three times independently. All statistical analysis were done using Microsoft excel or GraphPad Prism. *p*-values ≤0.05 were considered statistically significant as indicated by asterisks: *P≤0.05, **P≤0.01, and ***P≤0.001. For lifespan assay Log-Rank Test (Mantel-Cox Test) was performed. Asterix indicates ***P≤0.001.

## Results

### Generation of CG15117^-/-^ and CG15117^-/-^;CG2135^-/-^ double knockout strain

Functional β-glucuronidase (β-GUS) activity in Drosophila melanogaster is encoded by two genes, annotated as CG2135 and CG15117 (Bar et al., 2018). CG2135 was found to share 42.8% sequence identity and 61.5% similarity with human β-GUS, while 74% similarity and 62% identity with CG15117. In comparison, CG15117 and human β-GUS showed 64% similarity and 47% identity. Both *Drosophila* homologues conserve the key catalytic residues (Asp207, Arg382, His385, Glu451, Tyr504, and Glu540) and several MPS VII-associated mutations (e.g. L176, R216, R357, R382, R477, P408, A619) found in human β-GUS (Fig. 1) (Celik et al., 2021; Hassan et al., 2013; Islam et al., 1996, 1999; Jain et al., 1996; Oshima et al., 1987; Tomatsu et al., 2009; Vervoort et al., 1997). However, in vitro enzyme assays using purified enzyme revealed that CG2135 has a six-fold higher specific activity than CG15117. Accordingly, the CG2135 knockout model was established, which, despite reproducing cardinal features of MPS VII, retained ∼30% residual β-GUS activity due to CG15117 (Bar et al., 2018). To analyse whether the complete absence of β-GUS could aggravate the MPS VII-related phenotypes, we sought to generate a fly strain that lacks both CG15117 and CG2135.

**Figure 1:**
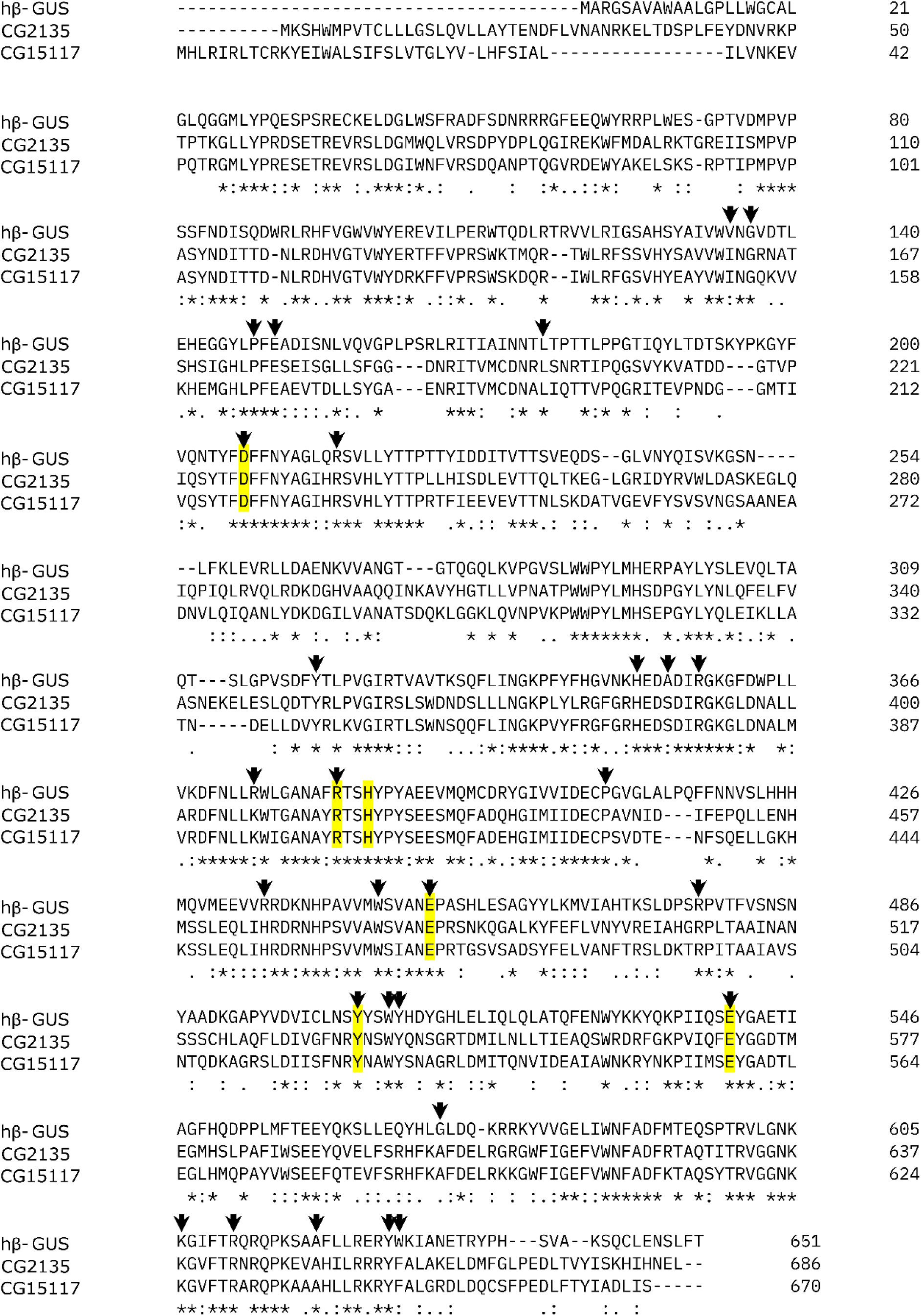
Multiple sequence alignment of human (hβ-GUS) and *Drosophila* β-Glucuronidases (CG2135 and CG15117). Multiple sequence alignment of amino acids from human (hβ-GUS) and *Drosophila* β-Glucuronidase enzymes, annotated as CG2135 and CG15117. Residues marked with an asterisk (*), colon (:), or period (.) denote fully conserved residues, residues with highly similar properties, or residues with less similar properties, respectively. Conserved Active site residues are highlighted in yellow. Black arrowheads denote conserved frequently mutated sites in the MPS VII patient.

To this end, we employed the CRISPR/Cas9 system to generate a CG15117 knockout (Port et al., 2014). Two gRNAs, targeting the 5′ and 3′ ends of CG15117, were designed using CHOPCHOP and validated by sequence alignment using BLASTn, which showed no off-targets (Labun et al., 2019). We employed both gRNAs to induce double-strand breaks at their target sites, followed by non-homologous end joining (NHEJ), which was expected to result in the deletion of an approximately 2.1 kb region encompassing all exons (Fig. 2A). To achieve this, the first requirement was to generate a transgenic fly expressing CG15117-specific sgRNAs. For this purpose, the selected gRNAs were cloned into a pCFD4 vector, which is predesigned to express two sgRNAs in *Drosophila* under the U6-1 and U6-3 promoter regions and has an attB site that facilitates its integration into the fly genome (Port et al., 2014). Transgenic flies expressing CG15117-specific sgRNAs were crossed with Nos-Cas9 flies (expressing Cas9 in the germ line cells) to induce germline deletions (Fig. S1). Progenies were screened for CG15117 knockout fly strains by genomic PCR with primers (P1-P2) flanking the target region expected to produce ∼1.8 kb amplicon (instead of 3.9 kb), confirming deletion of ∼2.1 kb spanning all exons (Fig. 2B). To confirm complete deletion of 2.1 kb region from CG15117 locus, internal primers (P3-P4) were used for genomic PCR, which produced no detectable product in the knockout lines (Fig. 2B). To confirm ablation of transcript, cDNA from CG15117^-/-^ flies was amplified using CG15117- specific primers, and no transcript was detected (Fig. S3A). Enzymatic assay of whole-fly lysates showed >50% reduction in β-GUS activity compared to wild-type (W1118), consistent with residual CG2135 expression. Interestingly, β-GUS activity levels in CG2135^-/-^ and CG15117^-/-^ lines were similar, suggesting comparable enzymatic contributions despite CG2135’s higher in vitro activity (Fig. S3B).

**Figure 2:**
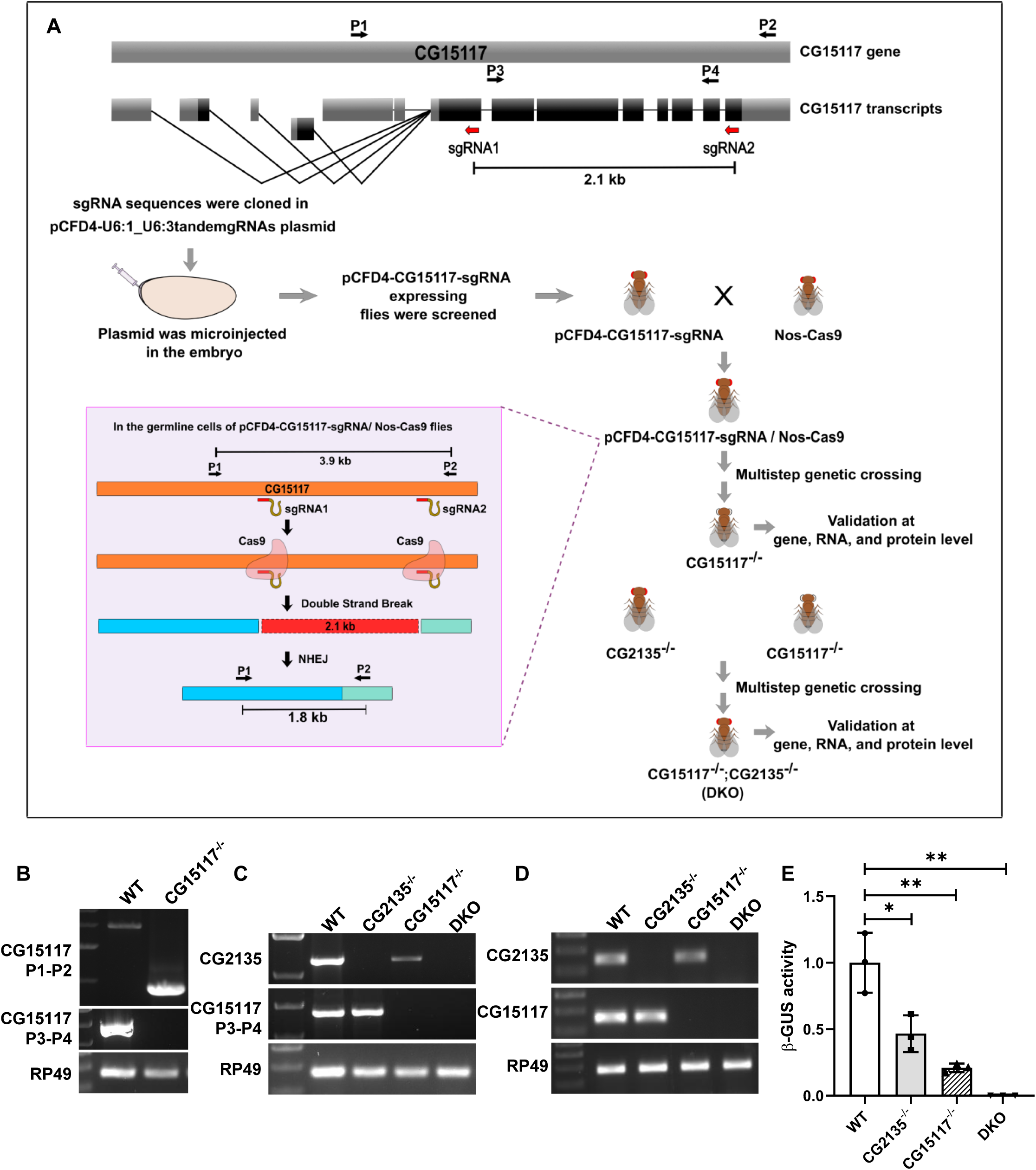
Generation and verification of CG15117^-/-^ and CG15117^-/-^;CG2135^-/-^ fly strains. (A) Graphical abstract showing the plan used to generate CG15117^-/-^ and CG15117^-/-^;CG2135^-/-^ double knockout fly strains. Schematic representation of the sgRNA target sites (red arrow) in the CG15117 gene for CRISPR-Cas9-mediated knockout and primer binding sites used for knockout screening. (B) Confirmation of the CG15117 knockout fly strain by genomic PCR using P1-P2 and P3-P4 sets. RP49 was used as a loading control. (C) Confirmation of CG15117^-/-^;CG2135^-/-^ double knockout (DKO) strains by genomic PCR using primers for CG15117 and CG2135. WT, CG2135^-/-^, CG15117^-/-^ were kept as control. (D) Confirmation of CG15117-/-;CG2135-/- (DKO) strains at transcript level PCR with CG15117 and CG2135 specific primers using cDNA as template. WT, CG2135^-/-^, CG15117^-/-^ were kept as control. (E) The bar graph represents the fold change in β-GUS activity in 4-day-old CG15117^-/-^, CG2135^-/-^, and DKO flies. The level of significance is shown with asterisks (*), *p≤0.05, **p≤0.01.

To generate a double knockout fly line that completely lacks β-GUS activity, CG2135^-/-^ and CG15117^-/-^ flies were crossed following the crossing scheme outlined in Fig. S2. The resulting CG15117^-/-^;CG2135^-/-^ (double-knockout) flies were validated for the absence of CG2135 and CG15117 at the genomic, transcript, and protein levels. For this, genomic DNA was isolated and used as a template to amplify CG15117 and CG2135 using specific primers, which shows the absence of CG2135 and CG15117 in the DKO fly genome (Fig. 2C). Similarly, the absence of the amplified product after PCR using specific primers for CG2135 and CG1517 and cDNA as a template proves the absence of these transcripts in CG15117^-/-^;CG2135^-/-^ (DKO) fly (Fig. 2D). We further validated the absence of CG2135 and CG15117 protein in DKO flies by the β-GUS activity assay using whole fly lysate. We observed complete absence of β-GUS activity validating the successful elimination of both gene products in the DKO strain (Fig. 2E).

### The effect of β-GUS depletion on survivability is greater in DKO flies

Our previous work demonstrated that CG2135 knockout (CG2135^-/-^) flies exhibit a reduced lifespan (Bar et al., 2018). Building upon this, we investigated the effects CG15117^-/-^ and double knockout (DKO: CG15117^-/-^;CG2135^-/-^) on longevity. For this, lifespan assays were performed for all three β- GUS-deficient genotypes (CG2135^-/-^, CG15117^-/-^, and DKO), alongside wild-type (WT) controls (Fig. 3A-B). The mean lifespan calculated from the survival graph was 60 days for WT, 59 days for CG15117^-/-^, 48 days for CG2135^-/-^, and 39 days for DKO (Fig. 3B). Similarly median lifespans were also obtained which revealed similar trends with median survival of 64 days for WT, 59 days for CG15117^-/-^, 50 days for CG2135^-/-^, and 39 days for DKO (Fig. 3B). Among all genotypes, DKO flies exhibited the shortest lifespan, with a significant reduction compared to WT, and a more pronounced decline than CG2135^-/-^ flies. These results emphasize the additive effect due to complete β-GUS loss. Interestingly, CG15117^-/-^ flies displayed no significant change in mean lifespan relative to WT flies. Collectively, these observations suggest that complete loss of β-GUS activity results in a greater impact on survival than loss of either gene alone. Increased susceptibility to starvation attributed to autophagy defect was one of the key observations in CG2135^-/-^ flies (Mandal et al., 2025). To further analyse the effect of partial vs complete loss of β-GUS activity on survivability under stress, we exposed all three β-GUS-deficient strains to starvation (complete and amino acid starvation). For this, 30 days old flies were subjected to complete starvation by transferring them to empty vials containing water-soaked tissue, and mortality was recorded every 12 hours. All three mutant strains showed increased sensitivity to starvation compared to age-matched WT flies. Among them, DKO flies were the most vulnerable, with a mean survival time (MST) of only 14.9 hours compared to the wild type (MST = 28.9 hours). Notably, CG15117^-/-^ flies also demonstrated increased susceptibility to starvation (MST = 23.46 hours), indicating a broader sensitivity to metabolic stress in the absence of β-GUS activity (Fig. 3C). To determine whether this effect was specific to total nutrient deprivation or extended to amino acid starvation, we further tested survivability on a 5% sucrose-only diet. In this assay, 30-day-old males from all genotypes were evaluated. DKO flies again exhibited significantly reduced survivability compared to WT. However, CG15117^-/-^ and CG2135^-/-^ flies did not show significant differences from WT under these conditions (Fig. 3D). These results suggest that complete β-GUS deficiency increases vulnerability to both total and amino acid starvation, while partial loss through single knockouts may not be sufficient to elicit this sensitivity under amino acid-limited conditions.

**Figure 3:**
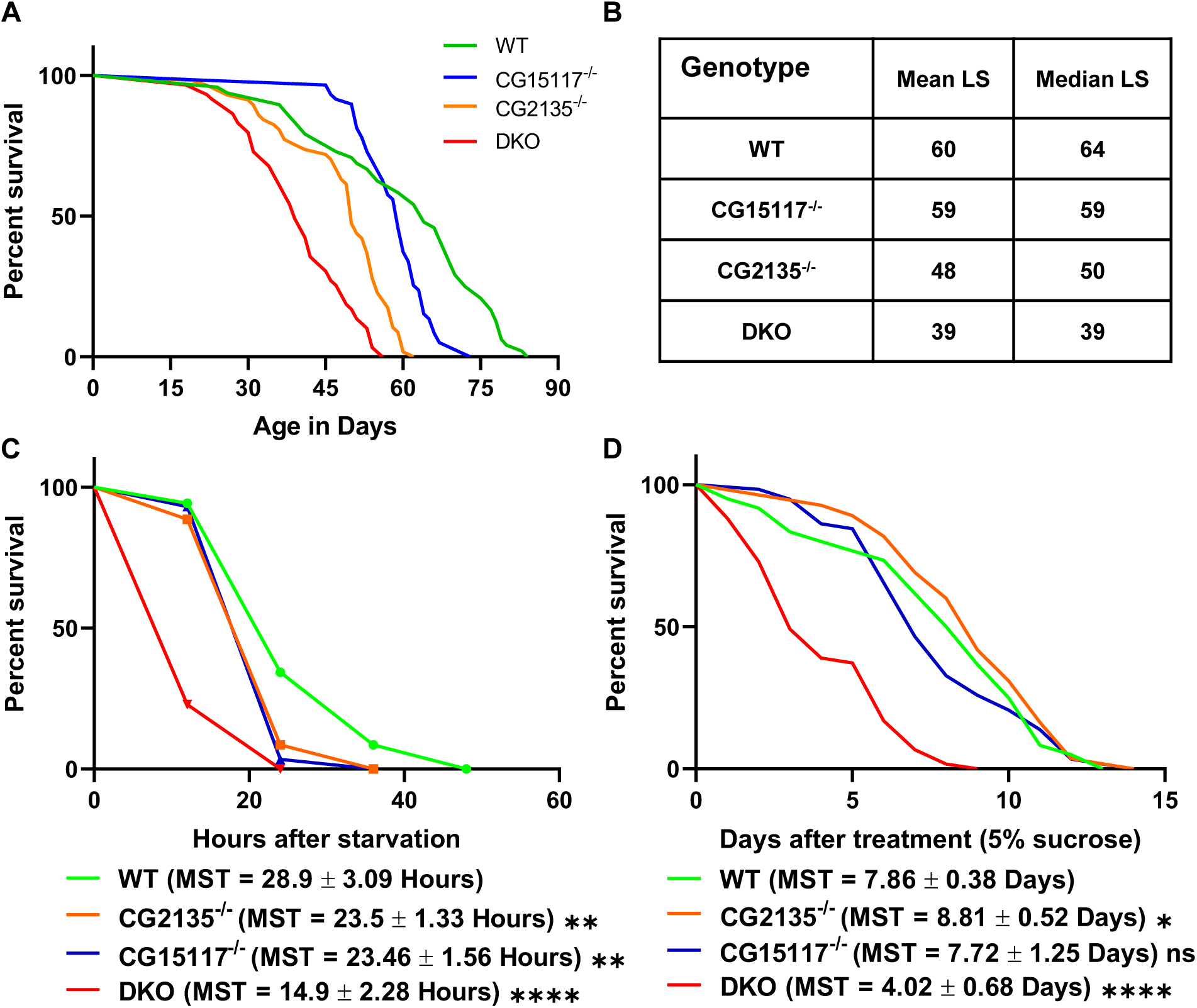
Comparative analysis of lifespan and susceptibility to starvation stress in β-GUS-deficient fly strains. (A) The line curve represents the percent survival of male wild type (WT), CG2135-/-, CG15117-/-, and DKO flies with age. N≥50 flies, Log-rank statistical test was performed, p>0.001. (B) The table shows the mean and median lifespan calculated for WT, CG15117-/-, CG2135-/-, and DKO flies from the survival curve (C). The line curve shows percent survivability of 30 day old β-GUS-deficient flies under amino acid starvation. Age matched WT flies were used as control group. N ≥ 50. Log-rank test was performed to check statistical significance. p<0.001. (D) The line curve shows percent survivability of 30 days old β-GUS-deficient under complete starvation. Age matched WT flies were used as control group. N ≥ 50. Log-rank test was performed to check statistical significance. p<0.001.

### Muscle ATP loss is the initial driver of locomotory defect in β-GUS-deficient flies

Loss of locomotor function is a well-characterized clinical manifestation in patients with Mucopolysaccharidosis VII (MPS VII) (Ballabio & Bonifacino, 2020; Zielonka et al., 2017). Consistent with this, our previous studies have shown that CG2135^-/-^ flies exhibit a progressive decline in climbing ability with age (Bar et al., 2018). To evaluate whether complete depletion of β-Glucuronidase (β-GUS) activity aggravates this phenotype, we conducted a systematic analysis of locomotor performance across β-GUS-deficient fly strains with age (Fig. 4A). Negative geotaxis assay (climbing assay) with 7-, 15-, 21-, 30-, and 45-day-old flies revealed progressive loss of climbing ability in CG2135^-/-^ and DKO fly strains with age compared to wild-type flies (Fig. 4B). Notably, DKO flies exhibited impaired climbing as early as day 7, which further deteriorated with age, dropping to less than 20% by day 30. In contrast, CG2135^-/-^ flies showed relatively normal performance until 21 days, and a 50% decline was observed when they reached 30 days of age. Interestingly, CG15117^-/-^ flies displayed no significant locomotor deficits compared to WT at any assessed time point, including at 45 days, when both CG2135^-/-^ and DKO flies exhibited near-complete loss of climbing ability (<5%) (Fig. 4B). Overall, these results demonstrate that the severity of locomotor decline correlates with the extent of β-GUS depletion. DKO flies exhibited the most profound impairment, followed by CG2135^-/-^ flies, whereas CG15117^-/-^ flies largely retained normal climbing ability. Interestingly, these fly strains mimicked varying degrees of disease severity observed in MPS VII patients, depending on the acquired mutation in the *β-GUS* gene. We recently demonstrated a correlation between locomotory impairment in CG2135^-/-^ flies and apoptotic neurodegeneration driven by ATP loss as a result of mitochondrial dysfunctionality in their brain. To understand if early locomotory loss in DKO flies was also due to elevated mitochondrial dysfunction, we analysed brain ATP levels in CG2135^-/-^ and DKO flies. As anticipated, we observed a marked 60% decrease in the ATP level in 45-day-old CG2135^-/-^ and DKO fly brains with increased mitotracker intensity in comparison to age-matched wild-type controls (Fig. S4A-H). Interestingly, to our surprise, we observed no change in the brain ATP level at 30 days old CG2135^-/-^ or DKO flies, despite a clear locomotory defect being observed (Fig. 4C). This temporal discrepancy suggests that the early onset of locomotor defects is unlikely to be driven by brain ATP depletion and may instead reflect dysfunction in other tissues. Muscle, being another crucial tissue involved in locomotion, we checked for ATP levels in 30-day-old CG2135^-/-^ and DKO muscles with age-matched wild-type flies as controls. Our analysis showed more than 90% reduction in ATP levels in CG2135^-/-^ and DKO flies compared to the wild type flies, indicating mitochondrial dysfunction in muscles precedes brain (Fig. 4D). Whether severe loss in the ATP levels leads to muscle degeneration in these flies at this early stage, we applied TUNEL staining to detect apoptotic nuclei in CG2135^-/-^ and DKO muscles. Our study revealed elevated apoptotic cells in these fly muscles compared to age-matched wild-type flies, indicating prominent muscular degeneration at this stage (Fig. 4E-H). These observations clearly indicate a temporal variation in the onset of mitochondrial dysfunction, with muscle being affected prior to brain tissue. Moreover, the locomotory defects observed in these β- glucuronidase-deficient flies were primarily triggered by muscle degeneration resulting from ATP depletion, and were subsequently exacerbated by additive neurodegeneration driven by an energy crisis.

**Figure 4:**
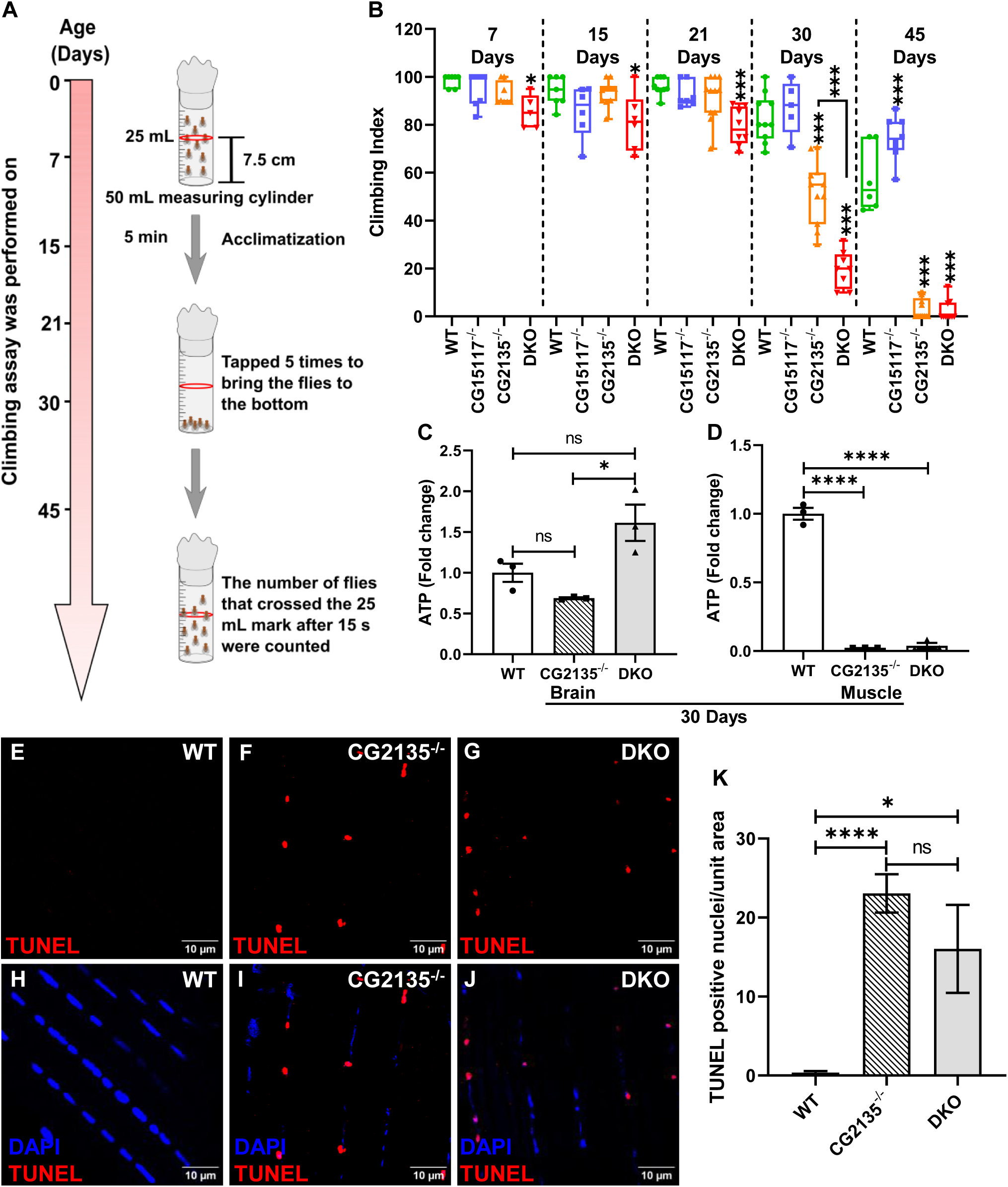
Progressive loss of climbing ability in CG2135-/- and DKO flies as a result of muscle ATP loss. (A) Schematic representation of the climbing assay, which was performed on 7-day, 15-day, 21-day, 30-day, and 45-day-old flies to detect locomotory defects. (B) The box plot with jittered points shows the climbing index of male β-GUS-deficient flies (CG15117^-/-^, CG2135^-/-^, and DKO) of 7-day, 15-day, 21-day, 30-day, and 45-day-old flies. Age-matched wild-type (WT) flies were treated as controls. N ≥ 100. (C) Bar graph showing fold change in ATP level in β-GUS-deficient flies compared to WT of 45- day-old fly brain (N=3). (D) The bar graph represents the quantification of total ATP from 30-day-old fly muscles (N=3), showing a reduction in CG2135-/- and DKO muscles compared to wild-type control. (E-J) TUNEL staining of 30-day-old fly muscles shows increased TUNEL-positive nuclei in CG2135-/- and DKO compared to age-matched WT counterparts. (K) The bar graph represents quantification of TUNEL-positive nuclei in the 40-day-old WT, CG2135-/-, and DKO fly muscle per unit area (per unit = 10000 μm^2^). N=4 fly.

### Impaired autophagy-lysosome mediated clearance in CG2135^-/-^ and DKO fly muscle

In line with our findings, we investigated the underlying cause of mitochondrial dysfunction in the CG2135^-/-^ and DKO muscles. Immunostaining of muscle revealed a significant increase in ATP5A signal in 30-day-old CG2135^-/-^ (∼2.72-fold) and DKO (∼2.20-fold) flies relative to age-matched wild-type controls (fig. 5A-G). Defective mitophagy has been shown to cause damaged mitochondrial accumulation in the 45-day-old CG2135^-/-^ fly brain (Mandal et al., 2025). With this prior knowledge, we checked the autophagy-lysosome axis in CG2135^-/-^ and DKO fly muscles as well. Our study revealed a reduced ATG8aII to ATG8aI ratio in 30-day-old CG2135^-/-^ and DKO fly muscles, indicating a defective autophagy turnover (Fig. 5H,J) (Klionsky et al., 2012). To further evaluate autophagy flux, we measured levels of Ref(2)P, the Drosophila homolog of the mammalian autophagy adaptor protein p62 (Nezis et al., 2008). Densitometric analysis of western blot data revealed a comparable >50% reduction in Ref(2)P levels in the muscles of the CG2135^-/-^ and DKO flies (Fig. 5H,K). Together, these findings provide strong evidence of defective autophagy-mediated clearance in the muscle tissue of CG2135^-/-^ and DKO flies, likely contributing to the progressive mitochondrial dysfunction and muscle pathology described in these models. However, comparable ATP loss and autophagy defect in CG2135^-/-^ and DKO fly muscle failed to explain strain-specific variations in locomotor dysfunction. We hypothesized that this discrepancy could be a result of variation in the degree of lysosomal dysfunction. To test this hypothesis, we examined the levels of mature Cathepsin L, a lysosomal protease commonly used as a marker of lysosomal activity. Elevated Cathepsin L activity has previously been reported in tissues from MPS VII animal models, including the mitral valve of affected dogs (Bigg et al., 2013; David et al., 2023). Western blot analysis of muscle lysates from 30-day-old flies revealed a pronounced increase in the mature form of Cathepsin L in DKO flies relative to wild-type, CG2135^-/-^ (Fig. 5H,I). These findings suggest that DKO flies experience more pronounced lysosomal dysfunction than either single knockout strain, potentially reflecting a more advanced pathological state analogous to severe forms of MPS VII. The elevation of mature Cathepsin L may represent a compensatory lysosomal response or an indicator of lysosomal overload and stress, further linking β-GUS deficiency to progressive muscle degeneration through disrupted lysosomal homeostasis.

**Figure 5:**
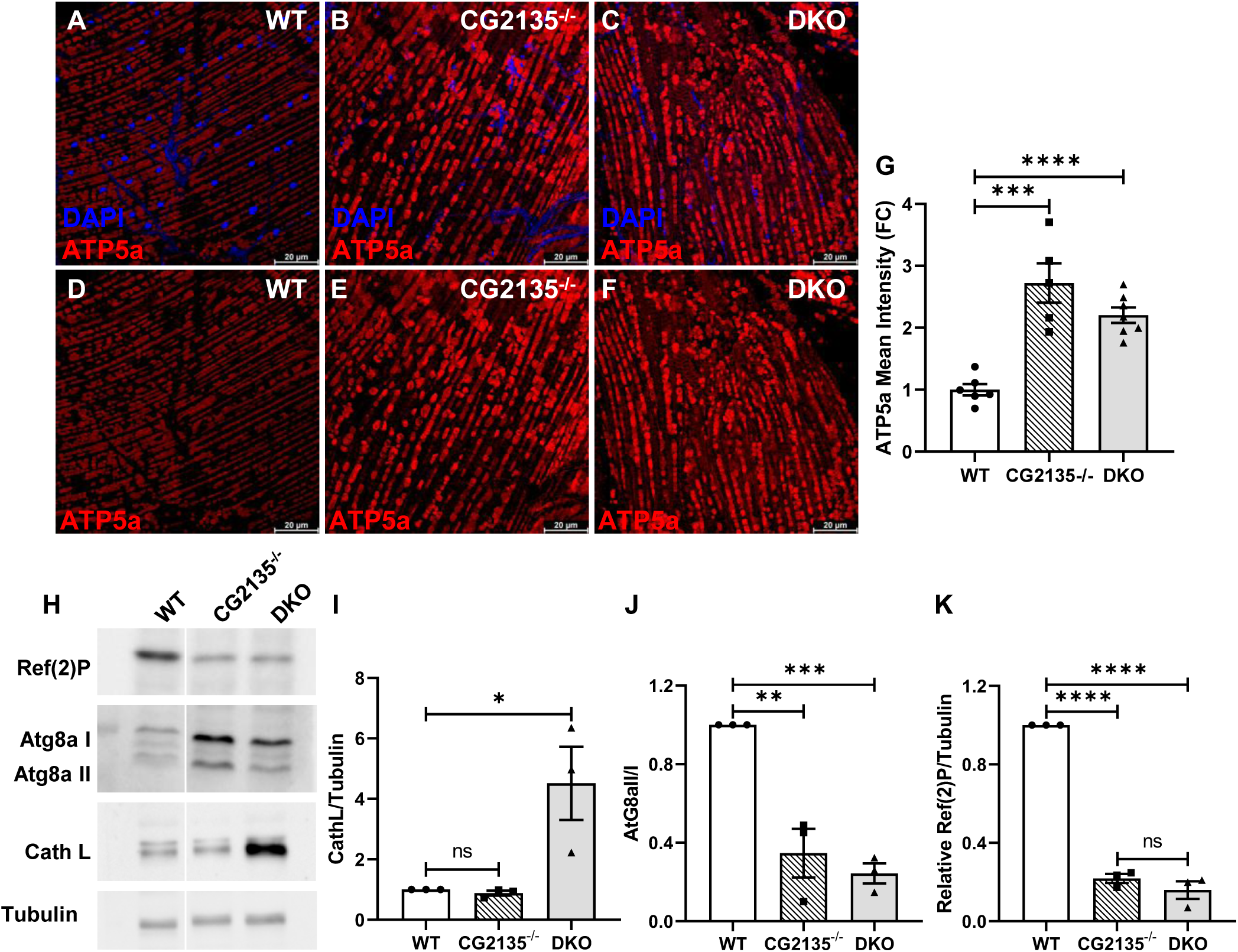
Accumulation of damaged mitochondria in 30-day-old CG2135^-/-^ and DKO fly muscle due to defective clearance. (A-F) Immunostaining of the thorax muscle of 30-day-old flies with ATP5a (red) to mark mitochondria and counterstaining with DAPI (blue) marks the muscle nuclei. (G) Bar graph showing quantification of the mean intensity of ATP5a (N≥5). (H) Representative western blot of Ref(2)P, Atg8aI-II, and Cathepsin L levels in the muscle of 30-day-old flies. Tubulin was used as a loading control. (I) The bar represents the quantification of the band intensity of Cathepsin L level normalized to the Tubulin level (N=3). (J) The bar represents the quantification of the ratio of band intensity of Atg8aII to that of Atg8aI (N=3). (K) The bar represents the quantification of the band intensity of Ref(2)P level normalized to the tubulin level (N=3). Error bars depict the SEM values. The level of significance is shown with asterisks (*), *p≤0.05, **p≤0.01, ***p≤ 0.001, ****p≤ 0.0001, and ‘ns’ means non-significant.

## Discussion

Here, we report that the extent of β-glucuronidase (β-GUS) deficiency dictates disease progression and severity in MPS VII. Using Drosophila β-GUS knockout models, we demonstrated that complete enzyme loss produces more severe pathologies and markedly accelerates locomotor decline compared to partial loss of β-GUS activity. Interestingly, mitochondrial dysfunction and energy failure in muscles of β-GUS-deficient flies preceded brain tissue, leading to muscular degeneration and locomotor defect. These observations not only highlight gene dosage-dependent effects of β-GUS loss but also underscore the significance of tissue-specific variation in disease progression.

The presence of two genes encoding β-GUS is a unique characteristic of *Drosophila melanogaster*. Both single knockout strains exhibited residual β-GUS activity attributed to the presence of CG2135 in the CG15117^-/-^ background and CG15117 in the CG2135^-/-^ background. In contrast to our previous observation that purified CG2135 exhibits six-fold higher activity compared to CG15117, the residual activity levels in both knockouts were comparable (Bar et al., 2018). This observation suggests a potential regulatory interdependency, possibly through a shared regulatory mechanism.

The deletion of both genes resulted in severe pathologies in adult flies compared to CG15117^-/-^ and CG2135^-/-^ flies. Double knockout flies have shown the highest reduction in mean lifespan, followed by CG2135 knockout flies. Whereas CG15117^-/-^ flies have shown no significant reduction in lifespan compared to wild-type flies. Moreover, the DKO flies displayed heightened sensitivity to starvation compared to CG2135^-/-^ and CG15117^-/-^. Increased sensitivity to starvation could reflect severe defects in lysosomal function (Ballabio & Bonifacino, 2020; Terman et al., 1999). Previous studies on *Drosophila* models of Gaucher disease have reported similar starvation susceptibility due to impaired clearance pathways (Kinghorn et al., 2016). While total starvation significantly reduced the survivability of all β-GUS-depleted flies, DKO flies, in particular, showed a marked sensitivity to amino acid starvation, reflecting a more severe form of the disease. More importantly, it is worth noting that depending on the type of mutation in the human β-GUS gene, disease severity and age of disease onset can vary (Gitzelmann et al., 1978; Sewell et al., 1982). Similarly, the differential loss of β-GUS in Drosophila produced phenotypic variation in disease severity and onset, reflecting gene dosage-dependent effects comparable to those arising from varying mutation types in humans. Collectively, these insights position β-GUS-deficient Drosophila strains as a robust model system for delineating the molecular basis of MPS VII progression and advancing the design of therapeutic interventions.

Our previous study on CG2135^-/-^ flies demonstrated that pronounced neuromuscular degeneration led to progressive climbing defect (Bar et al., 2018). A similar, though more severe, age-associated deterioration in climbing performance is also observed in double knockout (DKO) flies. Recently, we established that elevated mitochondrial dysfunction in the brain of CG2135^-/-^ flies is one of the primary contributors to neurodegeneration. Notably, restoration of mitochondrial function by induction of mitophagy was found to improve climbing ability in the CG2135^-/-^ fly (Mandal et al., 2025). In this study, we observed that mitochondrial dysfunction and ATP loss in CG2135^-/-^ and DKO fly brains were evident only at the very advanced stage of the disease. However, a significant climbing defect was already evident from an earlier age, specifically in DKO flies. This discrepancy raised the question of what might be triggering the locomotory defects. Further investigation to find the cause of locomotor decline revealed more than 90% reduction in total ATP level in the muscle of CG2135^-/-^ and DKO flies at a comparatively earlier age of 30 days. This drop in ATP level was also accompanied by elevated apoptosis, as evidenced by increased TUNEL-positive nuclei in muscles of these fly strains at this age. High-energy-demanding tissues like muscle are more vulnerable to ATP loss and often found to undergo apoptotic degeneration (X. Chen et al., 2023; Feldenberg et al., 1999; Lieberthal et al., 1998). Interestingly, no significant change was observed for ATP levels in the CG2135^-/-^ and DKO fly brains at this stage, indicating muscle degeneration as a plausible cause of the observed locomotory defect at this stage.

Muscle wasting has been associated with several lysosomal storage diseases, including MPS VII (Bar et al., 2018; Hesselink et al., 2003; Morishita & Petty, 2011). In mucopolysaccharidosis, fibroblasts are specifically vulnerable, as these are known to be one of the primary sites for GAG synthesis and secretion (Fuller et al., 2005; Merrilees et al., 1990). Notably, in milder forms of MPS VII, neurological abnormalities are either absent or manifest only at advanced stages, whereas musculoskeletal defects emerge earlier, indicating greater susceptibility to β-GUS deficiency (De Kremer et al., 1992; Montaño, Lock-hock, et al., 2016; Storch et al., 2003). However, the mechanistic events leading to muscle degeneration in MPS VII remained poorly explored. Our observation revealed autophagy defects in the muscles of 30-day-old β-GUS-depleted flies, adding a preliminary insight into mechanistic understanding. In CG2135^-/-^ fly brains, defective autophagosome biogenesis was indicated by reduced Atg1 transcript levels and decreased Atg8aII and Ref(2)P levels (S. Chen et al., 2014; Lin et al., 2024; Liu et al., 2023). Similarly, 30-day-old CG2135^-/-^ and DKO fly muscles also exhibited reduced Atg8aII/I and Ref(2)P levels, indicating autophagy impairment as a systemic feature of β-GUS deficiency.

In most lysosomal storage diseases, research has focused on the end-stage pathological outcomes, with limited attention to the early events driving disease progression. Our study offers preliminary insight into the mechanistic cascade underlying neuromuscular degeneration in β-GUS-deficient *Drosophila*. Our study showed that the muscle degeneration manifests as locomotor defects, which worsen in later stages as neuronal death becomes more apparent. We also observed a stepwise increase in phenotype severity corresponding to the differential loss of β-GUS activity, from single knockouts (CG15117^-/-^ or CG2135^-/-^) to double knockouts. These observations in particular suggests that differences in energy demand, GAG load, or clearance capacity might be the reason for temporally stratified organ vulnerability. In conclusion, our study revealed novel insight linking defective autophagy, mitochondrial dysfunction, and energy failure as the underlying causes of muscle degeneration in β- GUS-deficient *Drosophila*. Preceding mitochondrial dysfunction in muscle tissue than brain provides a novel insight into tissue-specific vulnerability due to loss of β-GUS activity. Collectively, our findings suggest that muscle degeneration, driven by defective autophagy, mitochondrial dysfunction, and severe energy crisis, underlies the onset of locomotor defects in β-GUS-deficient Drosophila.

## Acknowledgements

The authors express their sincere gratitude to Mohit Prasad for his support in fly maintenance and for providing critical reagents. The authors also thank Subhankar Dolai for his constructive comments on the work and Subhajit Majumdar for technical assistance. The support of the microinjection facility at cCAMP is gratefully acknowledged.

## Author contributions

N.M. and R.D. conceptualized the work and designed the experiments; N.M. and S.B. performed the experiments; N.M. S.B. and R.D. analysed the data; N.M. and R.D. wrote the manuscript; R.D. supervised the work and acquired the funding.

## Supplementary Materials

**Supplementary Figure 1:**
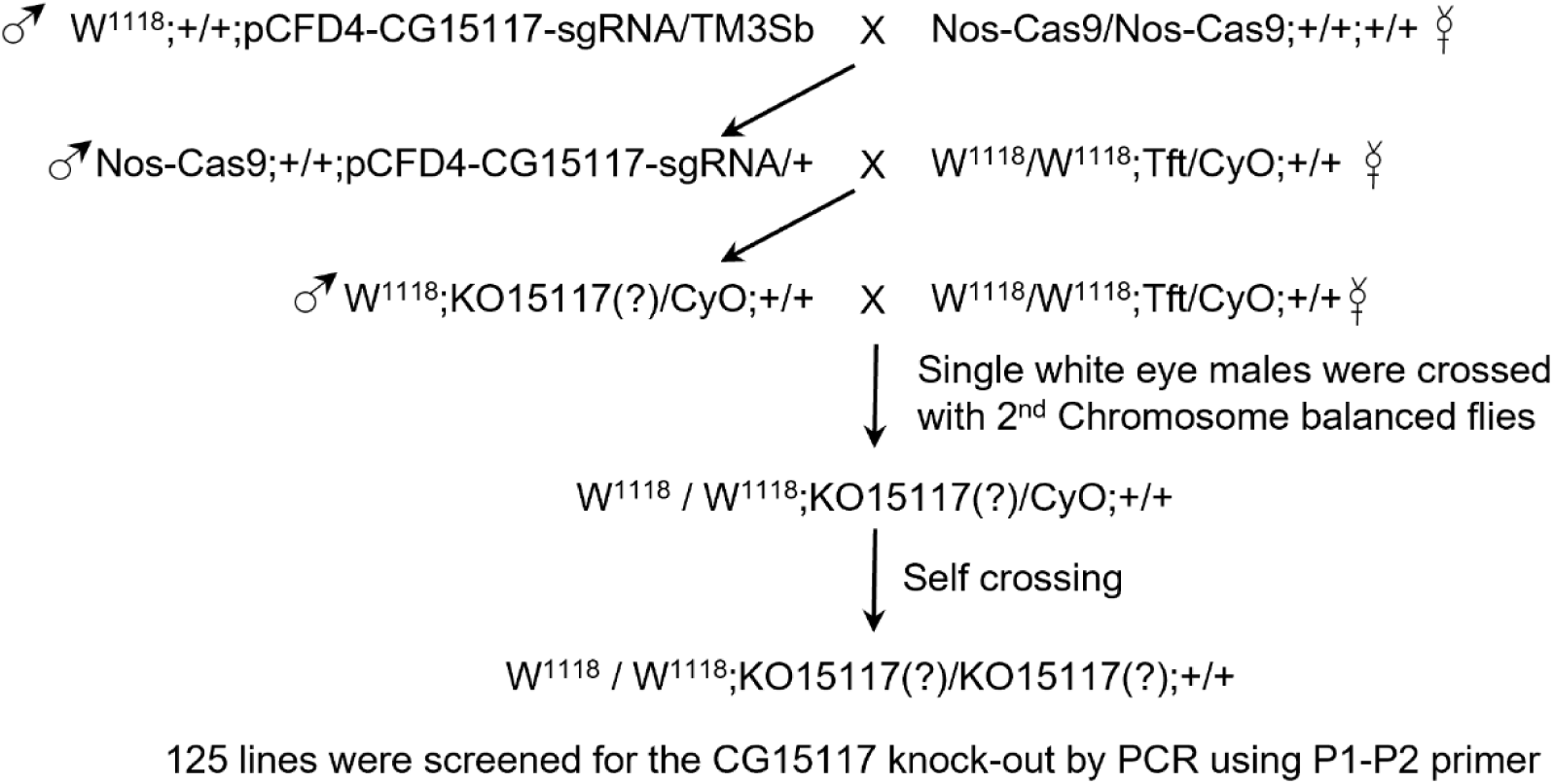
Crossing scheme used to generate CG15117^-/-^ knockout strain.

**Supplementary Figure 2:**
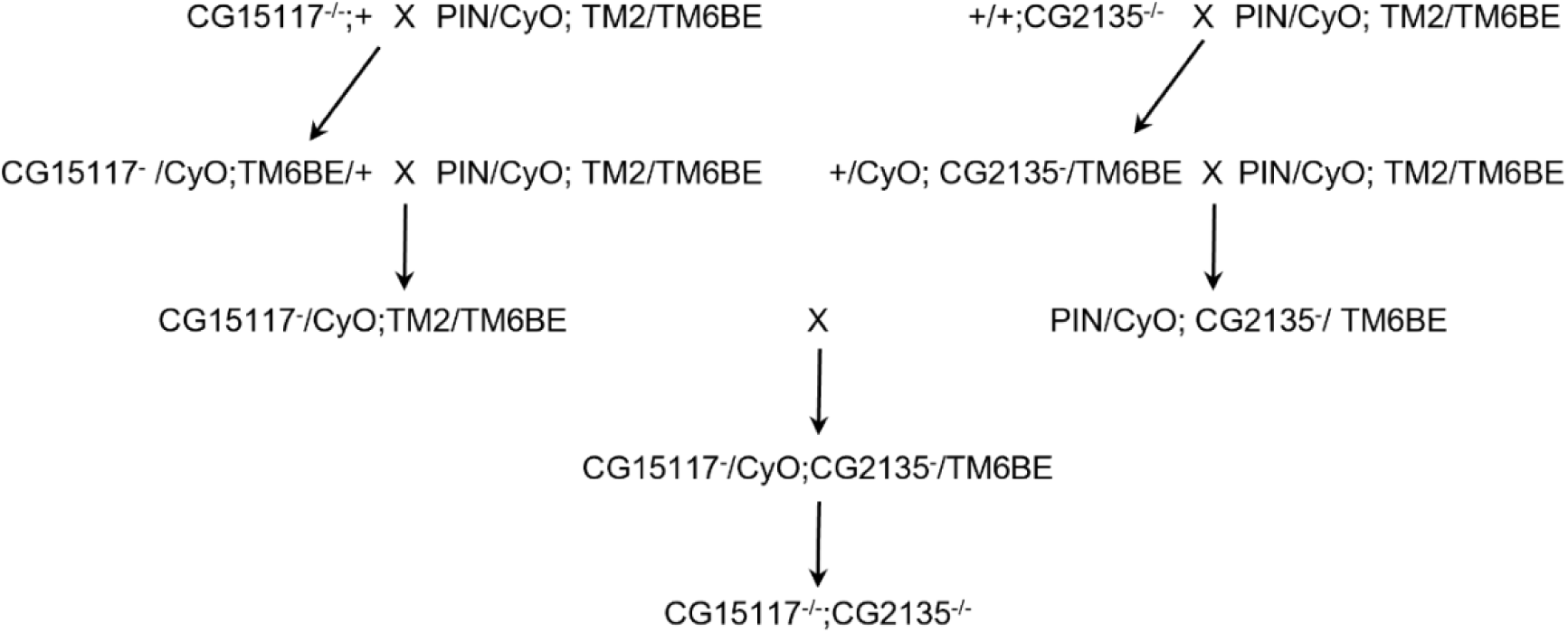
Crossing scheme used to generate CG15117^-/-^;CG2135^-/-^ double knockout strain.

**Supplementary Figure 3:**
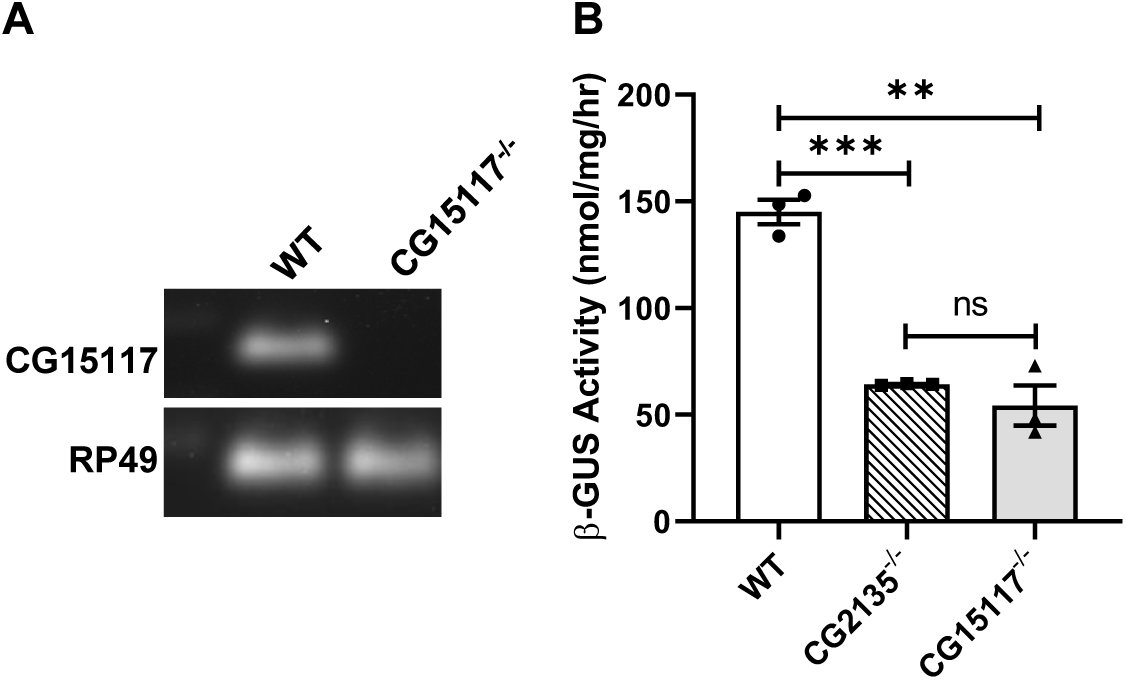
Generation and confirmation of CG15117^-/-^ fly. (A) Confirmation of CG15117^-/-^ knockout at transcript level by PCR with CG15117-specific primers using cDNA as template. (B) The bar graph represents specific activity of β-GUS in 4-day-old WT and CG15117^-/-^ flies, showing depletion of β-GUS activity in CG15117^-/-^ compared to matched WT control. Error bars depict the SEM values of three independent experiments. The level of significance is shown with asterisks (*), **p≤0.01, ***p≤0.001, and ‘ns’ means non-significant.

**Supplementary Figure 4:**
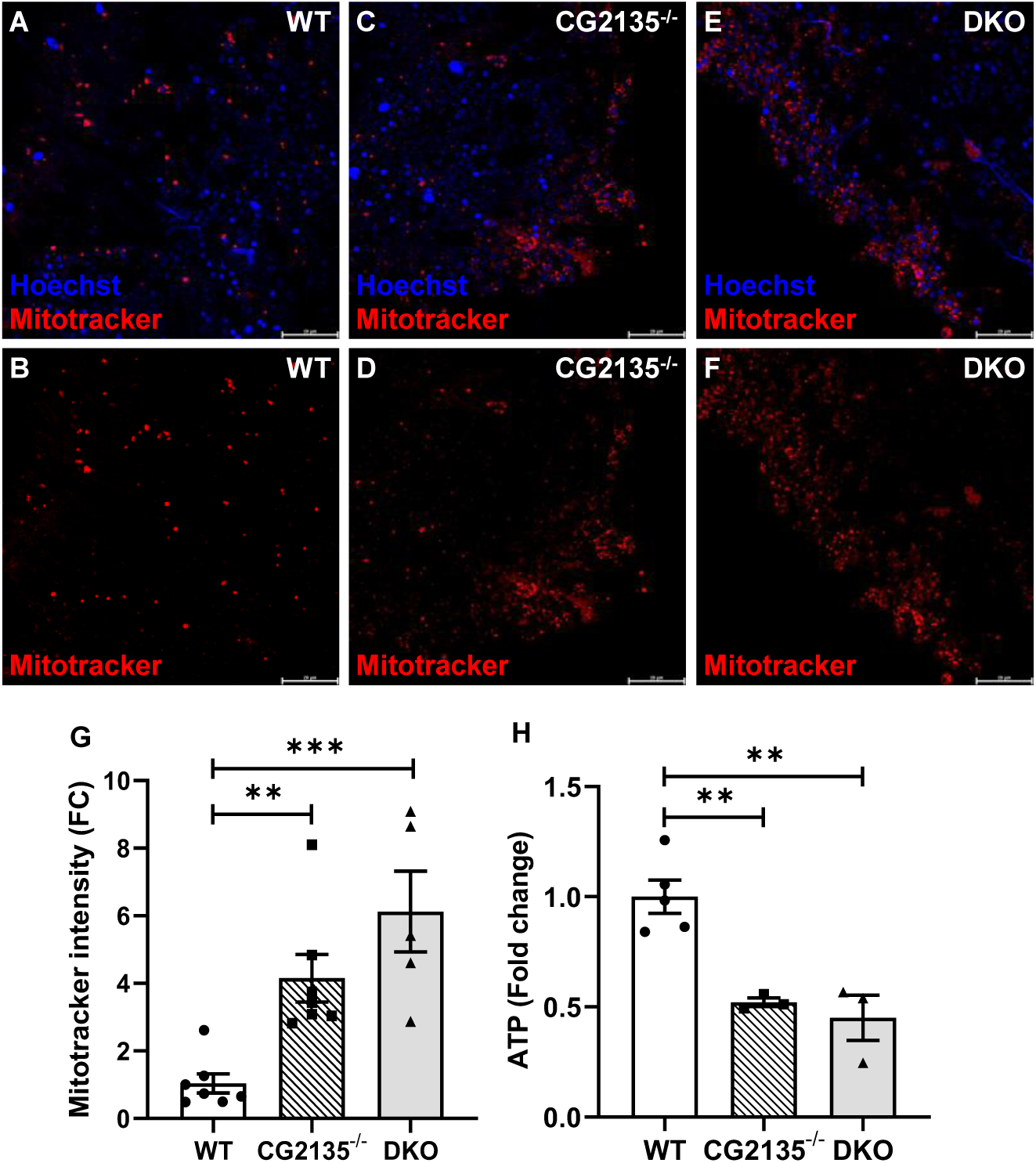
Mitochondrial defect is prominent in 45 days old CG2135^-/-^ and DKO fly brain. (A-F) Optic lobes of wild type (WT), CG2135^-/-^, CG15117^-/-^, and DKO fly brain (45 days old) stained for mitochondria with mitotracker red (in red) and nucleus with Hoechst (in blue), indicating accumulation of mitochondria in CG2135^-/-^ and DKO fly brain. (G) Quantification of mitotracker intensity revealed increased mitochondrial mass in these fly brains at 45 days compared to WT controls (N ≥ 5 fly brains). (H) The bar graph represents the quantification of total ATP from 45-day-old fly brains (N = 3), showing reduction in CG2135^-/-^ and DKO brains. Error bars depict the SEM values. The level of significance is shown with asterisks (*), **p≤0.01, ***p≤0.001, and ****p≤0.0001.

**Supplementary Table 1:**
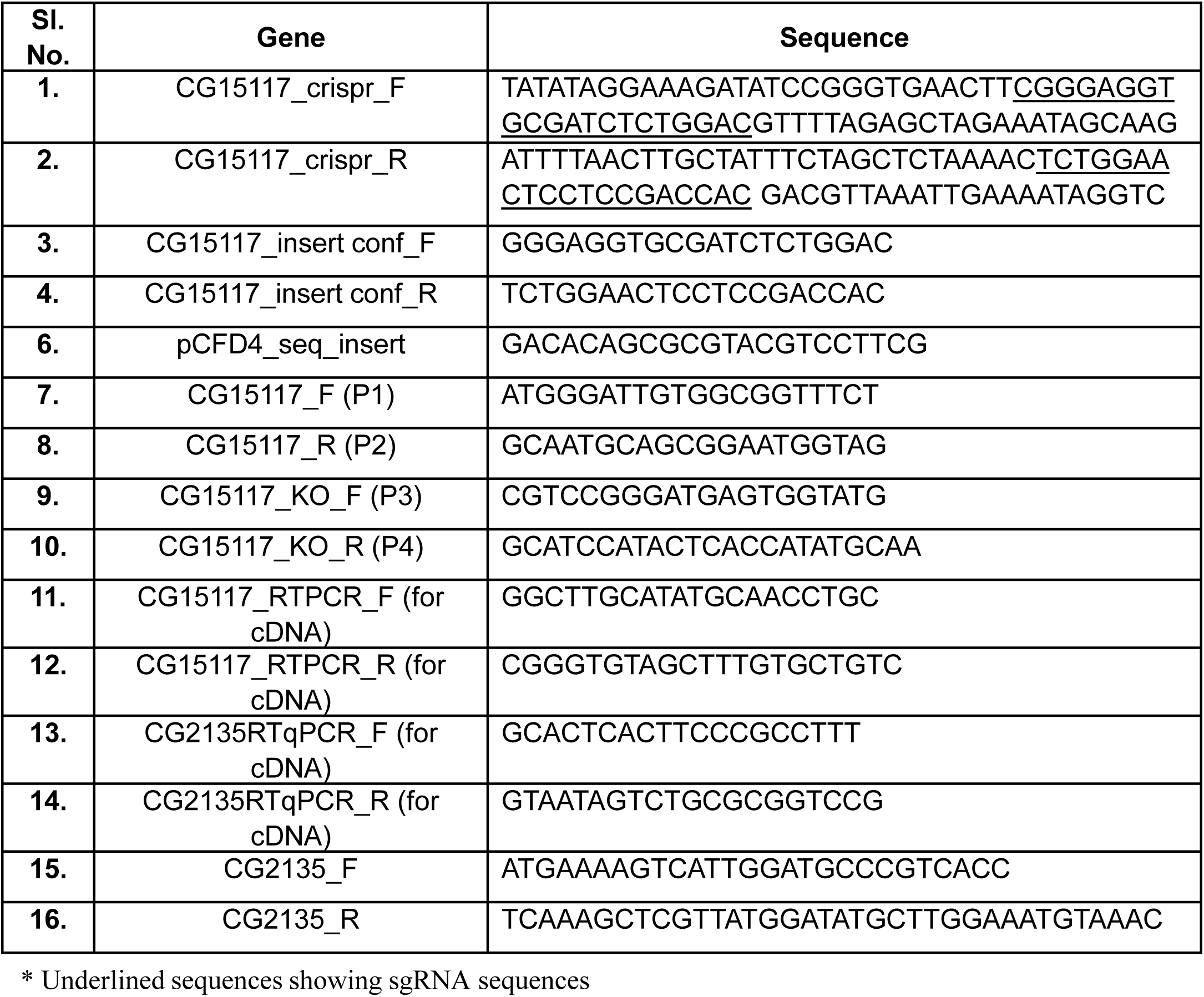
Oligonucleotides used in this study.

**Supplementary Table 2:**
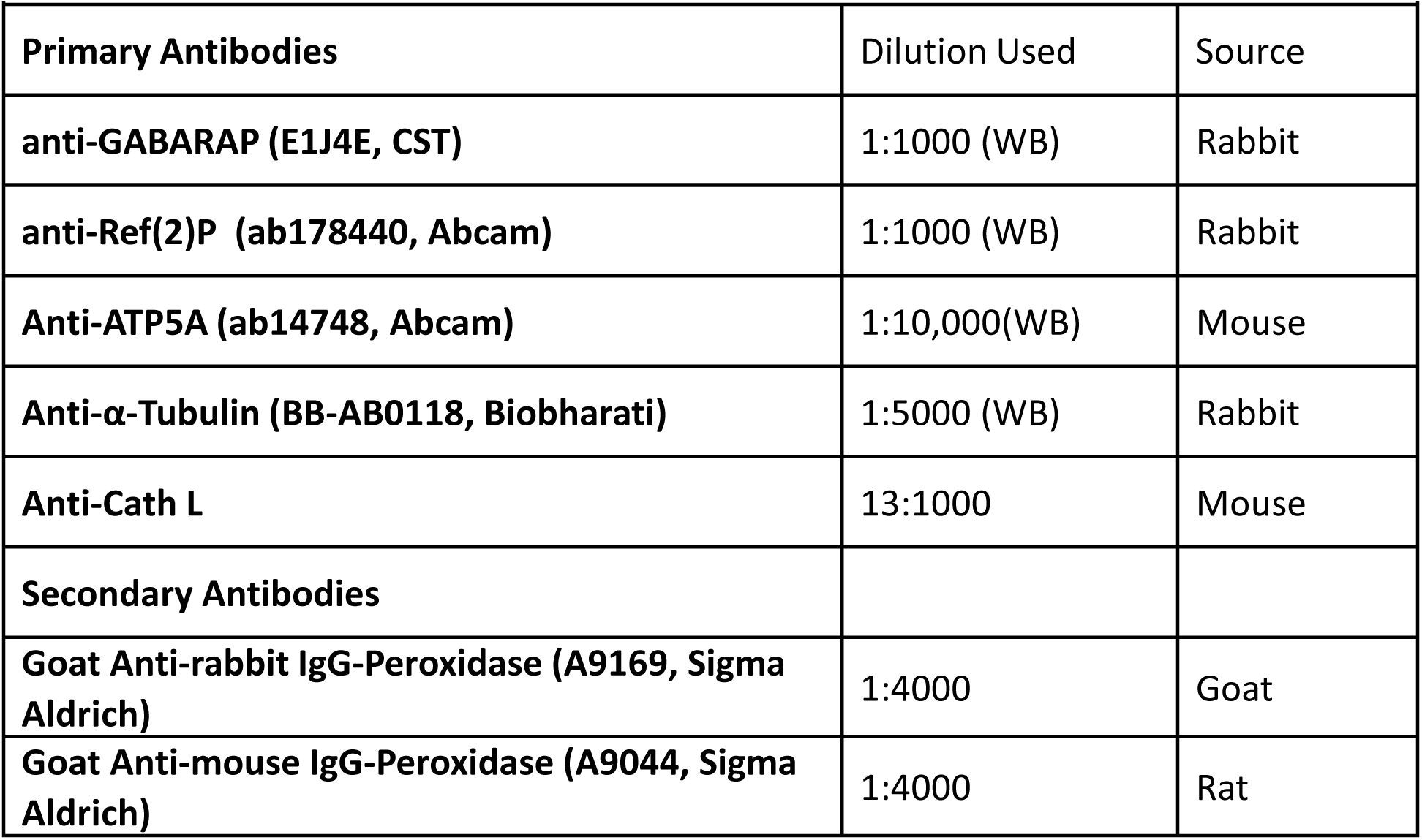
Antibodies used for Western blot.

| **Primary Antibodies** | **Dilution Used** | **Source** |
| --- | --- | --- |
| **anti-GABARAP (E1J4E, CST)** | 1:1000 (WB) | Rabbit |
| **anti-Ref(2)P (ab178440, Abcam)** | 1:1000 (WB) | Rabbit |
| **Anti-ATP5A (ab14748, Abcam)** | 1:10,000(WB) | Mouse |
| **Anti-α-Tubulin (BB-AB0118, Biobharati)** | 1:5000 (WB) | Rabbit |
| **Anti-Cath L** | 13:1000 | Mouse |
| **Secondary Antibodies** |  |  |
| **Goat Anti-rabbit IgG-Peroxidase (A9169, Sigma Aldrich)** | 1:4000 | Goat |
| **Goat Anti-mouse IgG-Peroxidase (A9044, Sigma Aldrich)** | 1:4000 | Rat |

